# Polyamine biosynthesis in *Xenopus laevis:* the gene xlAZIN2/xlODC2 encodes a lysine decarboxylase

**DOI:** 10.1101/661843

**Authors:** Ana Lambertos, Rafael Peñafiel

**Author notes:** Correspondence (RP).

## Abstract

Ornithine decarboxylase (ODC) is a key enzyme in the biosynthesis of polyamines, organic cations that are implicated in many cellular processes. The enzyme is regulated at the post-translational level by an unusual system that includes antizymes (AZs) and antizyme inhibitors (AZINs). Most studies on this complex regulatory mechanism have been focused on human and rodent cells, showing that AZINs (AZIN1 and AZIN2) are homologues of ODC but devoid of enzymatic activity. Little is known about *Xenopus* ODC and its paralogues, in spite of the relevance of *Xenopus* as a model organism for biomedical research. We have used the information existing in different genomic databases to compare the functional properties of the amphibian ODC1, AZIN1 and AZIN2/ODC2, by means of transient transfection experiments of HEK293T cells. Whereas the properties of xlODC1 and xlAZIN1 were similar to those reported for their mammalian orthologues, xlAZIN2/xlODC2 showed important differences with respect to human and mouse AZIN2. xlAZIN2 did not behave as an antizyme inhibitor, but it rather acts as an authentic decarboxylase forming cadaverine, due to its affinity for L-lysine as substrate; so, in accordance with this, it should be named as lysine decarboxylase (LDC). In addition, AZ1 stimulated the degradation of xlAZIN2 by the proteasome, but the removal of the 21 amino acid C-terminal tail, with a sequence quite different to that of mouse or human ODC, made the protein resistant to degradation. Collectively, our results indicate that in *Xenopus* there is only one antizyme inhibitor (xlAZIN1) and two decarboxylases, xlODC1 and xlLDC, with clear preferences for L-ornithine and L-lysine, respectively.

## Introduction

Ornithine decarboxylase (ODC) is a rate-limiting enzyme in the polyamine biosynthetic pathway that catalyzes the formation of putrescine from L-ornithine [1]. The polyamines spermidine and spermine, and their precursor putrescine, are organic cations that interact with different macromolecules, such as nucleic acids and proteins, affecting numerous cellular mechanisms related to cell growth and differentiation, signal transduction, apoptosis and autophagy [2–8]. In mammalian cells, ODC is highly regulated by a series of transcriptional, translational and post-translational mechanisms [1, 9–11]. Interestingly, ODC is a short-lived protein, with a half-live of less than 60 min in most mammalian tissues, and one of the few proteins that are degraded by the proteasome without ubiquitination [12, 13]. In addition, in the degradation of mammalian ODC, the antizyme 1 (AZ1) plays an important role [9, 14–16]. This regulatory protein is induced by increased levels of polyamines through an unusual ribosomal frame-shifting mechanism in the translation of AZ1 mRNA [17, 18]. AZ1 binds to the ODC monomer preventing the formation of the active ODC homodimer, and accelerates the proteasomal degradation of ODC, presumably by inducing the exposure of a cryptic proteasome-interacting surface of ODC [19]. The effects of antizymes on ODC are neutralized by antizyme inhibitors (AZINs), protein homologues of ODC but lacking decarboxylase activity [20–22]. In mammals, two AZINs have been identified (AZIN1 and AZIN2) that differ in their tissue expression profile [22–25]. In contrast to ODC, the degradation of these proteins is ubiquitin-dependent and is decreased by binding to AZ1 [26, 27].

Most studies on the structure, function and expression of ODC, AZs and AZINs have been carried out with the human and rodent versions of these proteins, and in less extension with the yeast and protozoan orthologues [28–31]. *Xenopus laevis* and *Xenopus tropicalis* are clawed frogs that have been used as model organisms in developmental biology. However, little is known about polyamine metabolism in these two species, and most of these studies have been focused on the changes in ODC activity and polyamine levels during *Xenopus laevis* oogenesis [32–34]. By screening a cDNA library from *Xenopus laevis* eggs, a cDNA corresponding to ODC (XLODC1) was isolated and sequenced [35]. Later, a new paralogue of ODC from *Xenopus laevis* (named xODC2) was identified, and the study of its temporal and spatial expression pattern during early embryogenesis showed that this is quite different from that of xlODCl [36]. In addition, whereas transfection studies of ODC-deficient mutant C55.7 CHO cells with XLODC1 showed that the *Xenopus* enzyme was functional in this heterologous cellular model [33], to our knowledge, no data on the activity and properties of xODC2 are available. In the Ensembl and Xenbase genome databases three *Xenopus* ODC paralogues are annotated: ODC1, AZIN1 and AZIN2. xAZIN2 gene is also named as xODC2, but it is unclear whether the corresponding protein functions as an antizyme inhibitor or alternatively it is an authentic ornithine decarboxylase. Due to the remarkable properties of mouse AZIN2 found in our previous studies [37–42], it appears relevant to analyze the characteristics of its amphibian orthologue to determine whether this protein functions as an enzyme or as an antizyme inhibitor. In the present work, we have transfected HEK293T cells with expression vectors containing the ORF corresponding to xAZIN2, xODC1, and xAZIN1, and the enzymatic activities and polyamine levels of these transfected cells were compared with those transfected with their murine counterparts. We also analyzed the degradation of the *Xenopus* ODC homologues and the effect of AZ1 on this process. Our results indicate that in *Xenopus laevis*, in contrast to mammalian cells, there are two different decarboxylases of ornithine and lysine, and only one protein acting as an antizyme inhibitor.

## Materials and methods

### Materials

L-[1-^14^C] ornithine and L-[1-^14^C] lysine were purchased from American Radiolabeled Chemicals Inc. (St. Louis, MO, USA). Anti-Flag M2 monoclonal antibody peroxidase conjugate (A8592), goat Anti-Rabbit IgG antibody peroxidase conjugated (AP132P), EDTA, Igepal CA-630, cycloheximide, L-lysine, putrescine dihydrochloride, cadaverine dihydrochloride, spermidine trihydrochloride, spermine tetrahydrochloride, 1,6-hexanodiamine, 1,7-diaminoheptane, dansyl chloride, proteasome inhibitor MG-132 and protease inhibitor cocktail (containing 4-(2-aminoethyl)benzenesulfonyl fluoride, EDTA, bestatin, E-64, leupeptin, aprotinin) were obtained from Sigma Aldrich (St. Louis, MO). Lipofectamine 2000 transfection reagent, Dulbecco’s Modified Eagle Medium (DMEM GlutaMAX), foetal bovine serum (FBS) and penicillin/streptomycin were purchased from Invitrogen (Carlsbad, CA). Pierce ECL PlusWestern Blotting Substrate was from ThermoScientific (IL, USA). Rabbit anti-ERK2 antibody (SC-154) was purchased from Santa Cruz Biotechnology (Texas, USA). The Anti-DYKDDDDK G1 Affinity Resin and the DYKDDDDK peptide were obtained from GenScript. D,L-alpha-difluoromethylornithine (DFMO) was provided by Dr. Patrick Woster (Medical University of South Carolina, Charleston, SC). Gene and protein sequences were obtained from Xenbase (http://www.xenbase.org/, RRID:SCR_003280) and Ensembl (www.ensembl.org) genome databases.

### Cell culture and transient transfections

Human embryonic kidney cells (HEK293T), obtained from ATCC, were cultured in DMEM (Dulbecco’s modified Eagle’s medium), supplemented with 10% (v/v) fetal bovine serum, 100 units/ml penicillin, and 100 μg/ml streptomycin, in a humidified incubator containing 5% CO_2_ at 37°C. Cells were grown to ~80% confluence and then were transiently transfected with Lipofectamine 2000 using 1.5 μl of reagent and 0.3 μg of plasmid per well (12-well plates). In co-transfection experiments, the mixtures contained equimolecular amounts of each construct.

The plasmid pcDNA3 without gene insertion was used as negative control. After 6 h of incubation, the transfection medium was removed, fresh complete medium was added, and cells were grown for additional 16 hours. Cells were collected and homogenized as described below, whereas the culture media was used for polyamine analysis. In some cases, xlAZIN2 and xlODC1 were purified from the cell extracts by affinity chromatography using an anti-Flag resin (GenScript) in accordance with the instructions of the supplier. All the constructs used in transient transfections were cloned into the expression vector pcDNA3.1. The Flag epitope DYKDDDDK was added to the N terminus of xlODC1, xlAZIN1, xlAZIN2, xlAZIN2ΔC, xlAZIN2-mAZIN2, mODC and mAZIN2 and to the C terminus of functional isoforms of murineAZ1, AZ2 and AZ3. All the clones were generated and purchased from GenScript, and sequenced before use.

### Western blot analysis

Transfected HEK293T cells were collected in phosphate buffered saline (PBS), pelleted, and lysed in solubilization buffer (50 mM Tris–HCl pH 8, 1% Igepal and 1 mM EDTA) with protease inhibitor cocktail (Sigma Aldrich). The cell lysate was centrifuged at 14,000×g for 20 min. Equal amounts of protein were separated in 10% SDS-PAGE. The resolved proteins were electroblotted to PVDF membranes, and the blots were blocked with 5% non-fat dry milk in PBS-T (Tween 0.1%) and incubated overnight at 4 °C with the anti-Flag antibody conjugated to peroxidase (1:10000). Immunoreactive bands were detected by using ECL Plus Western Blotting Substrate. ERK2, detected by a rabbit anti-ERK2 antibody (Santa Cruz, USA), was used as loading control. Densitometric analysis was achieved with ImageJ software.

### Enzymatic measurements

Transfected HEK293T cells were collected in phosphate buffered saline (PBS), pelleted and lysed in solubilization buffer (50mMTris-HCl, 1% Igepal and 1mM EDTA). After centrifugation at 14,000 ×g for 20 min, 5μl of the supernatant were taken to a final volume of 50μl with buffer containing 10 mM Tris-HCl, 0.25M sucrose, 0.1 mM pyridoxal phosphate, 0. 2 mM EDTA and 1 mM dithiothreitol. Decarboxylating activity was assayed in the supernatant by measuring ^14^CO_2_ released from L-[1-^14^C] ornithine or L-[1-^14^C] lysine. The reaction was performed in glass tubes with tightly closed rubber stopper, hanging from the stoppers two disks of filter paper wetted in 0.5 M benzethonium hydroxide, dissolved in methanol. The samples were incubated at 37°C from 15 to 120 minutes, and the reaction was stopped by adding 0.5 ml of 2 M citric acid. The filter paper disks were transferred to scintillation vials, and counted for radioactivity in liquid-scintillation fluid. In some cases, the enzyme activity was calculated by measuring by HPLC the rate of diamine formation (putrescine or cadaverine), after incubation of the cell extracts with different concentration of L-ornithine or L-lysine.

### Polyamine analysis

Both intracellular polyamines and polyamines generated in the culture media of the transfected cells were measured by HPLC. Transfected HEK293T cells were collected in phosphate buffered saline (PBS), pelleted, and the polyamines were extracted from the cells by treatment with 0.4M perchloric acid. The supernatant obtained after centrifugation at 10,000xg for 10 min was used for polyamine determination. For extracellular polyamine analysis, a fraction of the cell culture media was concentrated with a Speedvac Concentrator (Savant Instruments Inc. Farmingdale, NY, USA), and the resulting residue was resuspended in 0.4 M perchloric acid and processed as described above. Polyamines from the acid supernatant were dansylated according to a standard method [43]. Dansylated polyamines were separated by HPLC using a BondaPak C18 column (4.6 × 300 mm; Waters) and acetonitrile/water mixtures (running from 70:30 to 96:4 during 30 min of analysis) as mobile phase and at a flow rate of 1 ml/min. 1,6-Hexanediamine and 1,7-heptanediamine were used as internal standards. Detection of the derivatives was achieved using a Waters 420-AC fluorescence detector, with a 340 nm excitation filter and a 435 nm emission filter.

### Confocal microscopy

Cells grown on coverslips were transfected with xlAzin2, xlOdc1, xlAzin1, mAzin2 or mOdc constructs. Twenty-four hours after transfection, cells were fixed with 4% paraformaldehyde in PBS and permeabilized with 0.5% Igepal in PBS. For detection of Flag-labelled proteins, cells were incubated with an anti-Flag M2 monoclonal antibody (1:7.000), followed by an Alexa 488-conjugated secondary antibody (1:400). For the staining of nucleus, cells were loaded with DAPI (1:10000) for 5 minutes. Finally, samples were mounted by standard procedures, using a mounting medium from Dako (Carpinteria) and examined with a Leica True Confocal Scanner TCS-SP2 microscope.

### Statistical analysis

The data were analyzed by Student’s t-test for differences between means. P < 0.05 was considered as statistically significant.

## Results

### Comparative study of gene and protein structure of *Xenopus* AZIN2 with its paralogues and mammalian orthologues

According to the Xenbase genome browser, the gene structures of Azin2 described for *Xenopus tropicalis* (XB-GENE-6454420) and *Xenopus laevis* (XB-GENE-6493979) are similar. The comparison of protein sequences between *Xenopus tropicalis AZIN2* (xtAZIN2) (NP_001015993.2) and *Xenopus* laevisAZIN2 (xlAZIN2) (NP_001079692.1), by using the Clustal omega sequence alignment program, revealed a high homology (93.64%) (S1 Fig). Since our preliminary experiments showed that both proteins behave similarly, we selected *Xenopus laevis* for most experiments.

Next, we compared the gene structure of *Xenopus laevis* Odc paralogues with their respective murine and human orthologues. Fig 1 shows that the xlAzin2 gene, like mouse Azin2 (mAzin2) and human AZIN2 (hAZIN2), is formed by 11 exons (9 of them are coding exons), whereas xlOdc and xlAzin1 contain 12 exons (10 coding exons), similarly to their murine and human orthologues. The protein homology between the different orthologues of *Xenopus laevis* and mice was analyzed by using the Align Sequences Protein BLAST (NCBI), and the results are shown in Table 1. The sequence homology of xlAZIN2 with respect to xlODC1 or mODC was higher (65% and 63%, respectively) than that ofmAZIN2 (59%). In addition, sequence similarity of mODC to xlODC1 was higher (82%) than that of xlAZIN2 (65%). The lowest identity of xlAZIN2 was with xlAZIN1 (43%). These results indicate that although the genetic structure of xlAzin2 is close to its mammalian orthologues, its protein sequence is closer to that of *Xenopus* or mouse ODC proteins.

**Fig 1.**
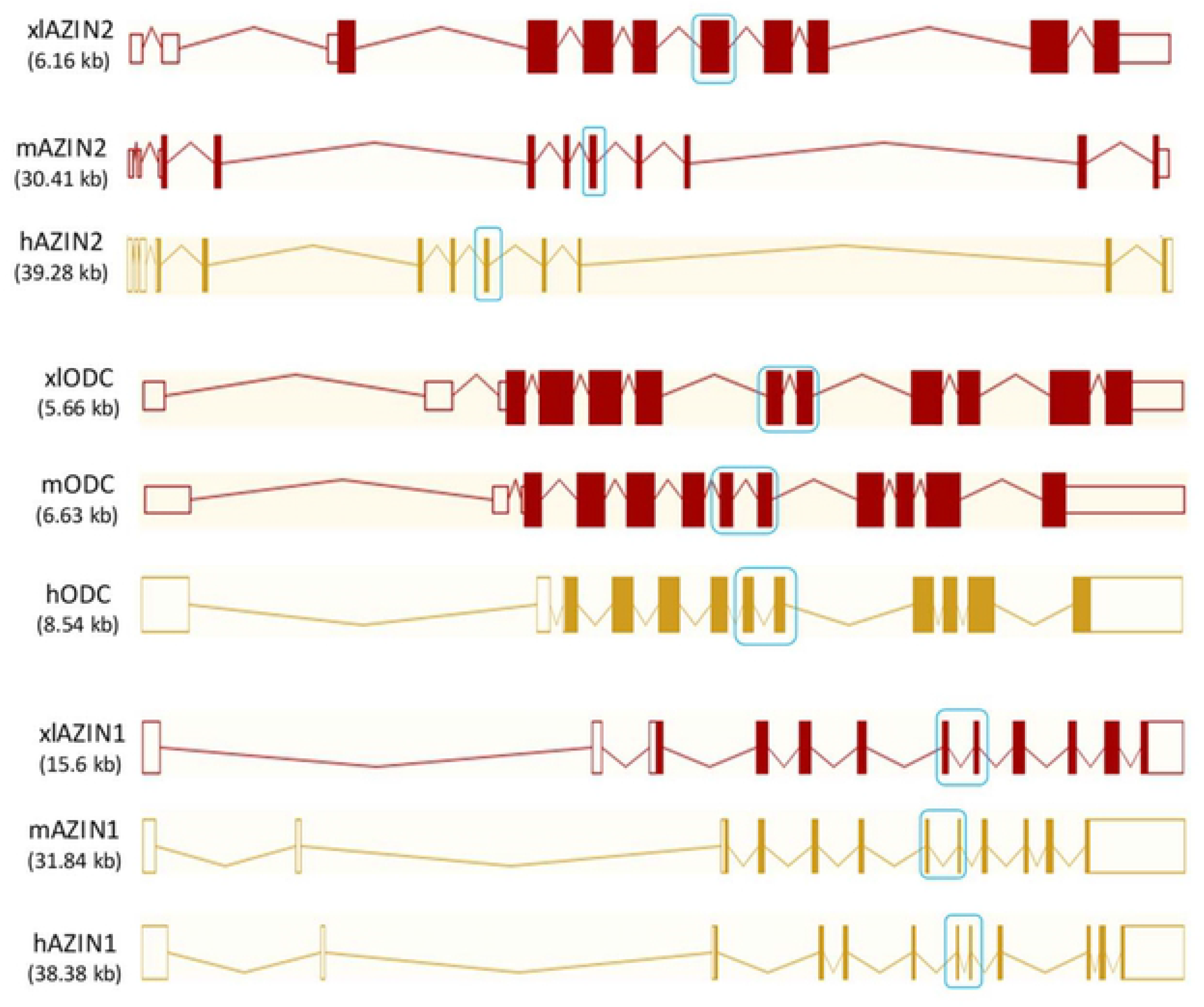
Genetic structure of mouse and human ODC paralogues, and their comparison with their *Xenopus* orthologues. Note that exons 7 and 8 in ODC and AZIN1 are fused in only one exon in AZIN2 (blue boxes). Data obtained from Ensembl (www.ensembl.org).

**Table 1.** Sequence identity between mouse (m) and *Xenopus laevis* (xl) homologue proteins.

|  | xlAZIN2 | xlODC1 | xlAZIN1 | mAZIN2 | mODC | mAZIN1 |
| --- | --- | --- | --- | --- | --- | --- |
| xlAZIN2 | 100 | 65 | 43 | 59 | 63 | 49 |
| xlODC1 | 65 | 100 | 47 | 52 | 82 | 51 |
| xlAZIN1 | 43 | 47 | 100 | 42 | 46 | 67 |

Fig 2 shows the sequence alignment of the proteins corresponding to the three ODC paralogues of *Xenopus laevis* (xlODC1, xlAZIN1 and xlAZIN2) and mODC. The amphibian xlAZIN2, as xlODC1, shares with mODC the 22 residues that are required for the decarboxylating activity [40, 44–48] whereas xlAZIN1, as reported for mammalian AZIN1 and AZIN2, lacks some essential residues such as K69 and C360. These results indicate that, according to these putative catalytic residues, xlAZIN2 appears to be closer to ODCs than to AZINs. Fig 2 also shows that lower homologies were found in the ~70 amino acids residues of the C-terminal region. The identity values of mODC with respect to xlODC1, xlAZIN2 and xlAZIN1 were 63%, 31% and 17%, respectively (S1 Table). Since two adjacent segments in the C-terminal region of ODC (segments S1 and S2 in S2 Fig), have been proposed as having different roles in the proteasomal degradation of ODC induced by AZ1 [19], we also calculated the sequence homology in these segments among the different ODC homologues (S2 Fig). S1 Table also shows that the identity values among S1 segments from mODC and its amphibian homologues (77%, 44% and 26%) were higher than those corresponding among the S2 segments (66%, 14% and 9%).

**Fig 2.**
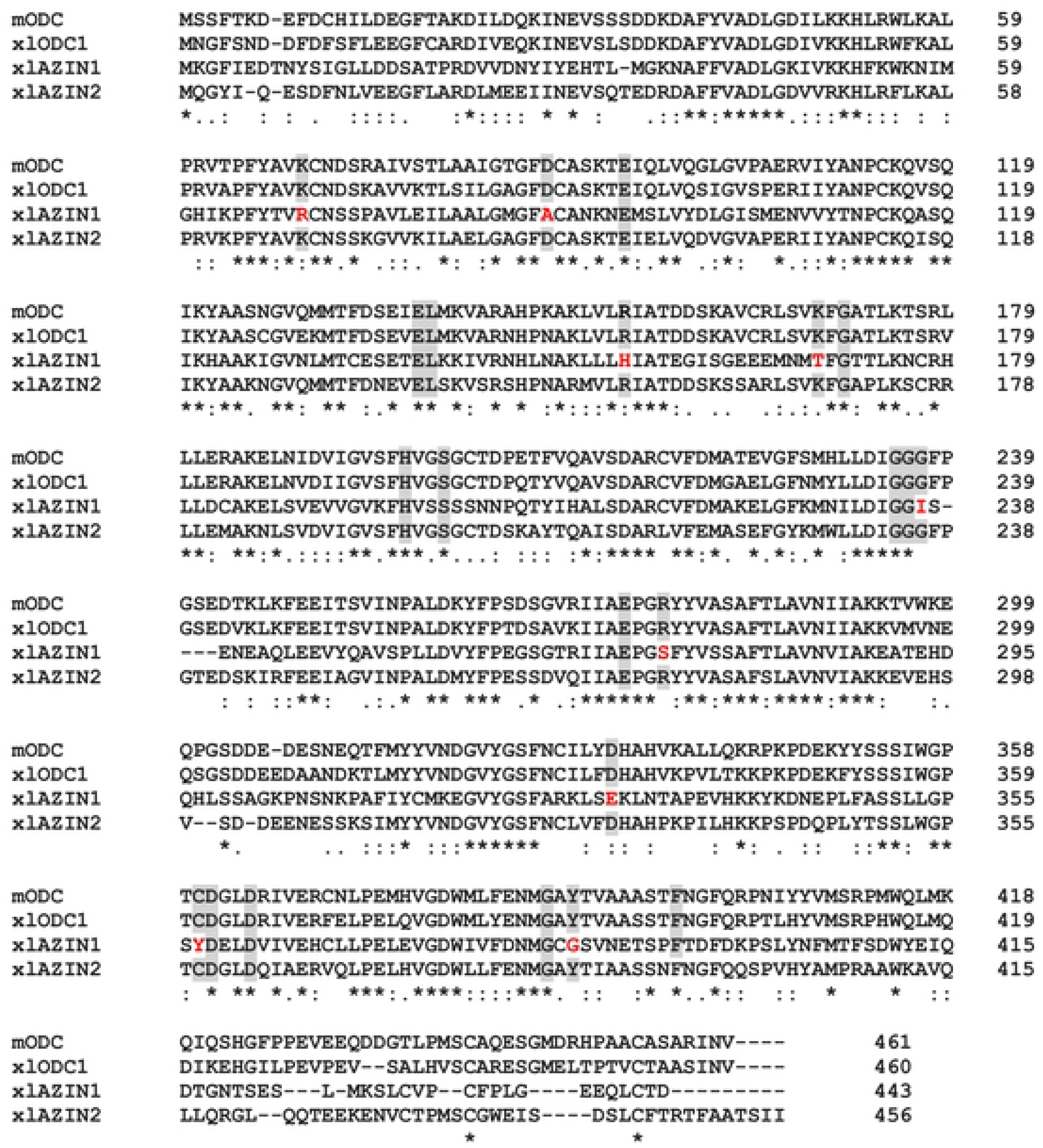
Comparison of the amino acid sequences of mouse ODC, xlODC1, xlAZIN1 and xlAZIN2 using ClustalW program for multiple sequence alignment. Asterisks represent amino acid identity; colon and dots represent amino acid similarity between the proteins. Grey background indicates amino acid residues associated with the catalytic activity of mODC that are conserved in the *Xenopus laevis* homologues. In red: substitutions in these critical residues in xlAZIN1.

### Functional analysis of xlAZIN2 in a heterologous cell system

To test the potential ornithine decarboxylase activity of xlAZIN2, HEK293T cells were transiently transfected with xlAZIN2, and the decarboxylating activity was measured in homogenates from the transfected cells. In parallel, cells were also transfected with the empty vector and with plasmids containing the coding sequences of xlODC1 and xlAZIN1, in the same vector as xlAZIN2. As displayed in Fig 3A, the homogenates from cells transfected with xlODC1 showed, as expected, a high ODC activity in comparison to those from mock transfected cells. In the case of xlAZIN2, the ODC activity was about 22% of the values found for xlODC1, and much higher than that of xlAZIN1. Western blot analysis revealed that these differences in ODC activity were not due to significant differences in protein expression levels. Both xlODC1 and xlAZIN2 were inhibited by treatment of the cells with 1 mM DFMO (Fig 3B). These results suggested that either xlAZIN2 is an antizyme inhibitor more potent than xlAZIN1 for increasing the endogenous ODC activity, or that it may possess intrinsic catalytic activity.

**Fig 3.**
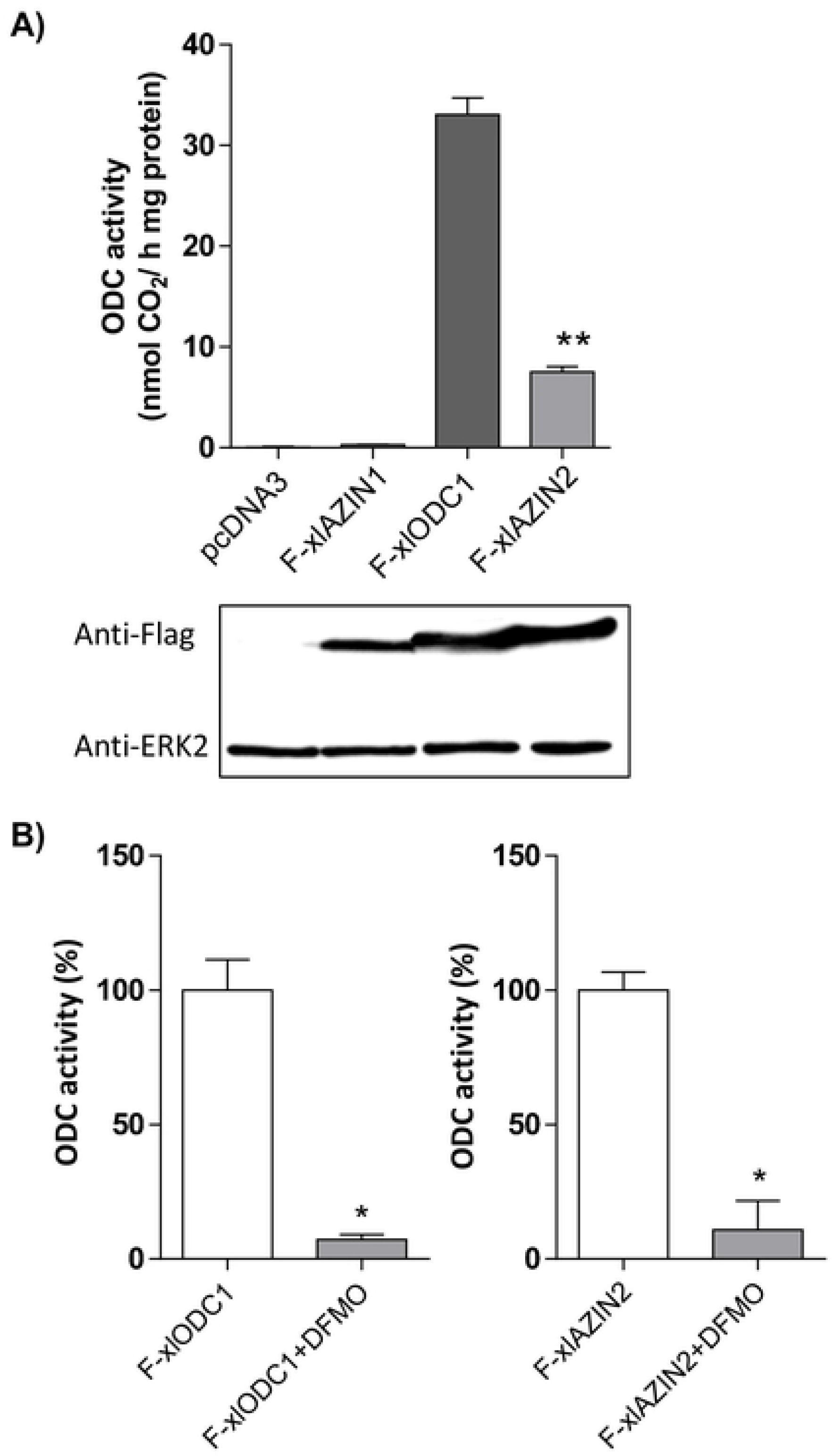
Expression of xlODC1, xlAZIN1 and xlAZIN2 in HEK293T transfected cells. HEK293T cells were transfected with the corresponding constructs of Flag-xlODC1, Flag-xlAZIN1, Flag-xlAZIN2 or empty vector, as indicated in the Experimental Procedures. (A) Top: ODC activity measured in the cell lysates. Bottom: Western blot analysis of the proteins detected using anti-Flag or anti-ERK2 antibodies. Results are expressed as mean±SE, and are representative of three experiments. (**) P<0.01 *vs* pcDNA3 or X-xlODC1. (B) Influence of 1mM alfa-difluoromethylornithine (DFMO) on the ornithine decarboxylase activity of xlODC1 and xlAZIN2 cell lysates. (*) P<0.05.

To corroborate the latter possibility, we next analyzed the influence of xlAZIN2 on polyamine levels. For that purpose, we studied the influence of xlAZIN2 transfection on the polyamine content of the transfected cells and on that of the culture media, after 16 h of the transfection.

Fig 4A shows the chromatogram traces of the dansylated polyamines obtained by HPLC analysis of HEK293T homogenates from cells transfected with xlAZIN2 or with the empty vector. A dramatic increase in putrescine levels was evident after transfection with xlAZIN2. Unexpectedly, the major increment was observed for cadaverine, the diamine that is produced by decarboxylation of L-lysine, with values about 3-fold higher than those of putrescine. The analysis of the polyamine content of the culture media of the xlAZIN2-transfected cells also showed that cadaverine was the most abundant polyamine, with values about 10-fold higher than those of putrescine (Fig 4B). The finding that the cadaverine to putrescine ratio in the cell cultures was about 3-fold higher than the diamine ratio in the cell extracts revealed that cadaverine is excreted more efficiently than putrescine in this type of cells.

**Fig 4.**
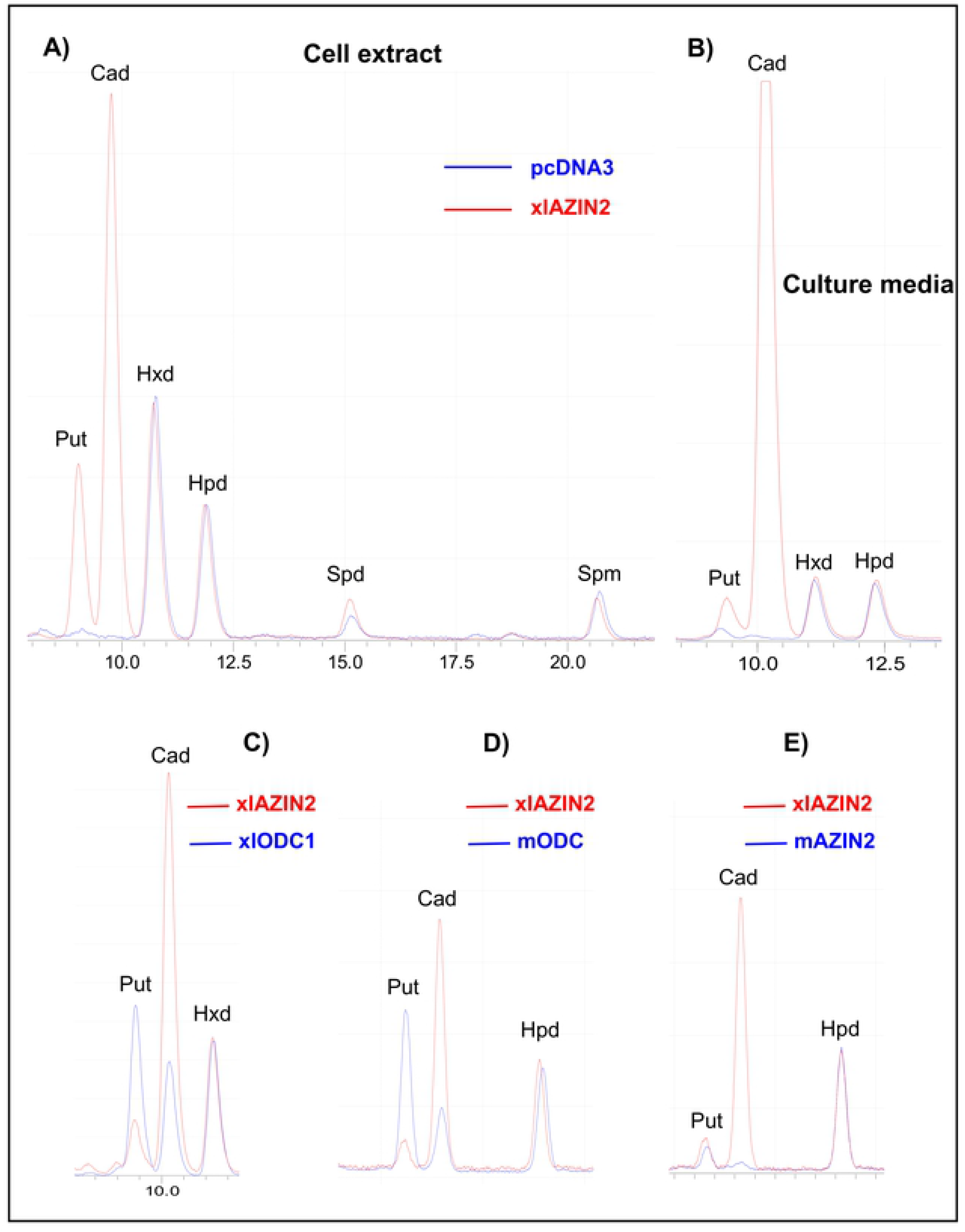
Analysis of the products formed by HEK293T cells transfected with different constructs. After 16 h of transfection, the culture media was aspirated and the cells collected. An aliquot of the media was concentrated and resuspended in perchloric acid 0.4 M, whereas the cells were homogenized in the same acid (200 μl per well). After centrifugation at 12.000 ×g for 15 min, the supernatants were dansylated and analyzed by HPLC as described in the Experimental section. (A) Overlapped HPLC chromatogram traces of the dansylated extracts from cells transfected with xlAZIN2 (red line) or with the empty vector pcDNA 3.1 (blue line). Hexanediamine (Hxd) and heptanediamine (Hpd) were used as internal standards. Put: putrescine; Cad: cadaverine; Spd: spermidine; Spm: spermine. (B) Overlapped HPLC chromatogram traces corresponding to the dansylated polyamines present in the culture media of cells transfected with xlAZIN2 (red line) or empty vector (blue line). (C) Comparison of the polyamines found in the culture media of cells transfected with xlAZIN2 (red line) with those of xlODC1, mODC and mAZIN2 (blue line).

Since it is known that mouse and rat ODCs are able to decarboxylate L-lysine, but less efficiently than L-ornithine [49], we compared the levels of putrescine and cadaverine in cells transfected with xlAZIN2 with those of the cells transfected with xlODC1, mODC or mAZIN2. Figs 4C and 4D show that the ratio cadaverine:putrescine in the cells transfected with any of the two ODCs were lower than one, whereas in the case of xlAZIN2 this ratio was higher than 7. These results indicate that xlAZIN2 is more efficient to synthesize cadaverine than putrescine under the cell culture conditions employed in the assays. In addition, in the cells transfected with mAZIN2, only vestigial levels of cadaverine were detected, whereas putrescine levels were similar to those of xlAZIN2 transfected cells (Fig 4E). All these results clearly indicated that xlAZIN2 behaves as an enzyme that can decarboxylate both amino acids L-ornithine and L-lysine to produce putrescine and cadaverine, respectively.

### Kinetic analysis of the decarboxylase activity of xlAZIN2

The enzyme kinetic parameters were analyzed using cell homogenates from xlAZIN2- or xlODC1-transfected HEK293T cells and different substrate concentrations. Table 2 shows that in the case of xlAZIN2 the Km for L-lysine (1.06±0.25 mM) was lower than the Km for L-ornithine (6.57±1.75 mM), suggesting that the affinity of xlAZIN2 to L-lysine is higher than the one to L-ornithine. The opposite was found for xlODC1, although here the affinity of xlODC1 for L-ornithine was much higher than for L-lysine (Km^Orn^=0.023±0.008mM and Km^Lys^=30.1±7.8 mM). Taking the ratio Vm/Km as an indicator of the catalytic efficiency of each enzyme, the results presented in Table 2 indicate that xlAZIN2 was much more efficient to decarboxylate L-lysine than xlODC1, whereas the opposite was found when L-ornithine was the substrate. In parallel experiments, using enzymes purified by affinity chromatography with anti-Flag beads, the Km values found were essentially similar to those presented in Table 2.

**Table 2.**
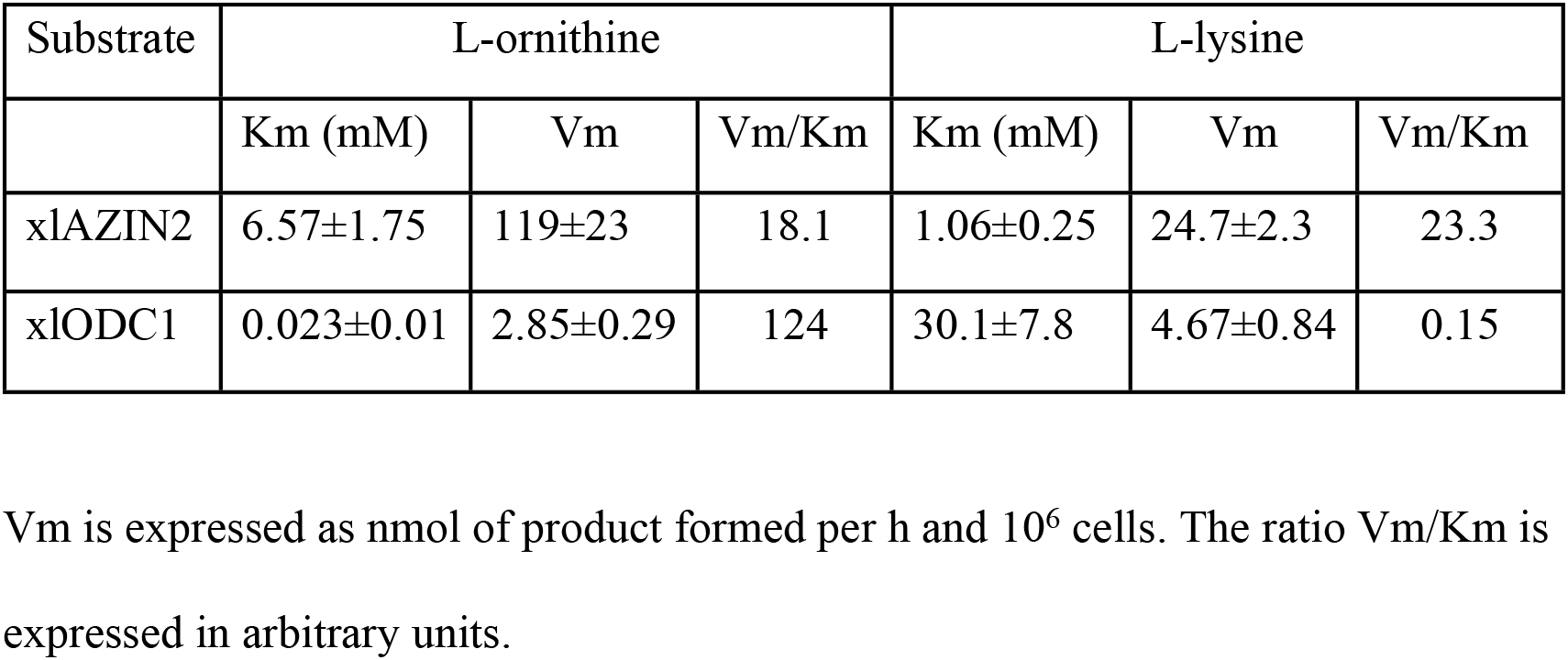
Comparison of the kinetic parameters of xlAZIN2 and xlODC1.

### Study of a possible antizyme inhibitory action of xlAZIN2

Although all above results clearly supported that xlAZIN2 has decarboxylating activity, it could be likely that xlAZIN2 may also act as an antizyme inhibitor. To test this possibility, we analyzed the ability of xlAZIN2 to rescue xlODC1from the predictable degradation induced by AZ1, as early reported for mouse AZIN2 [37]. To this purpose, we carried out different co-transfection experiments using several constructs. The results shown in Fig 5A corroborated that, as expected, AZ1 stimulated the degradation of xlODC1, and that none of the two xlAZIN2 constructs used (either with Flag for western blot detection or without Flag) were able to protect xlODC1 from degradation. In addition, the results shown in this figure also suggested that xlAZIN2 was induced to degradation by AZ1. To confirm this possibility, xlAZIN2 was co-transfected with AZ1, and the cell homogenates were analyzed for decarboxylase activity and xlAZIN2 protein content. Fig 5B clearly shows that AZ1 induced the degradation of xlAZIN2. Taking into consideration that earlier studies showed that mouse AZIN2 protected mouse ODC from degradation, whereas it was not degraded by AZ1 [37], the results shown here do not support a role of xlAZIN2 as an antizyme inhibitor. On the contrary, similar experiments using xlAZIN1 showed that AZ1, as it is known for mAZIN1 [26], protected the amphibian protein from degradation (S3 Fig).

**Fig 5.**
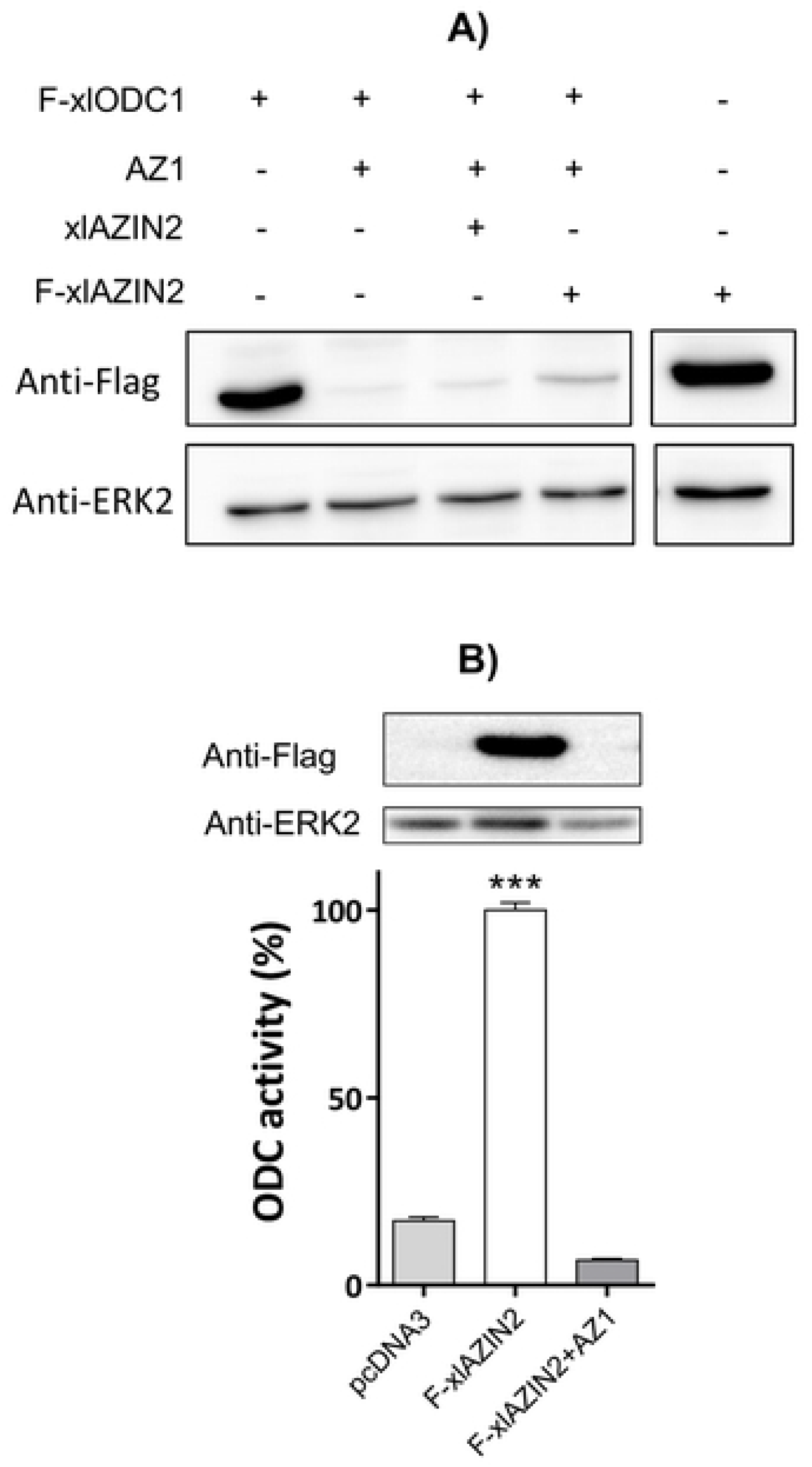
Influence of AZ1 on protein levels of xlODC1 and xlAZIN2. (A) Western blot of lysates of HEK293T cells co-transfected with xlODC1 and different combinations of AZ1 and xlAZIN2. (B) Western blot and ODC activity of lysates of cells co-transfected with Flag-xlAZIN2 and pcDNA3 or AZ1. (***) P<0.001 vs pcDNA3 or F-xlAZIN2+AZ1.

Furthermore, the subcellular distribution of xlAZIN2 in the transfected cells was found to be mainly cytosolic, similar to that of xlODC1 or mODC, and different from that of mAZIN2 (Fig 6). All these results demonstrate that the gene annotated as xlAZIN2 in the different gene databanks does not code for a *bona fide* antizyme inhibitor, but instead it encodes for an authentic amino acid decarboxylase with preference for L-lysine as substrate. Therefore, we propose to change its name to lysine decarboxylase (LDC).

**Fig 6.**
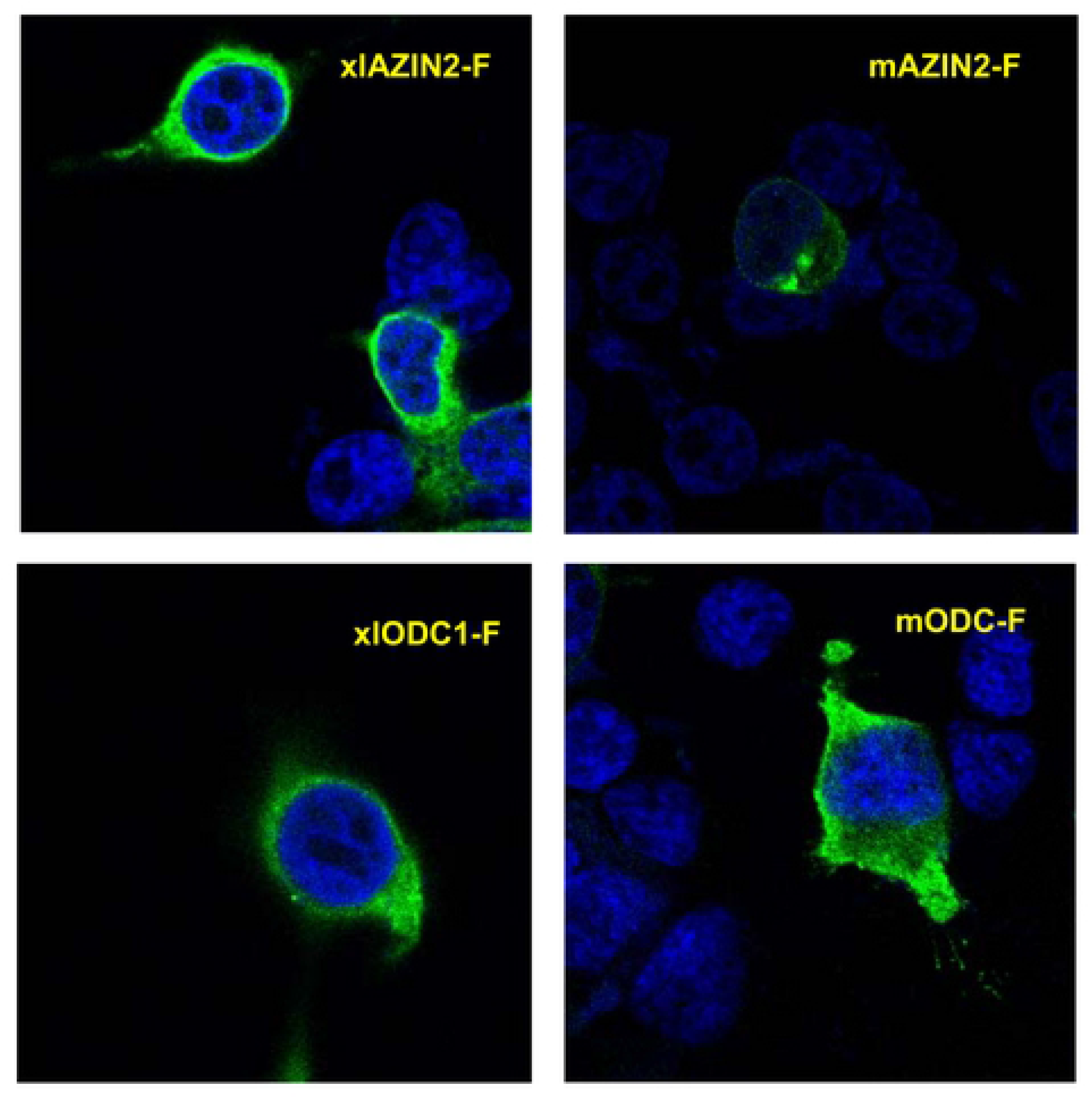
Subcellular location of xlODC1 and xlAZIN2 in transfected cells. Laser scanning confocal micrographs of HEK293T cells transfected with xlODC1, xlAZIN2, mODC or mAZIN2 fused to the Flag epitope. After transfections, cells were fixed, permeabilized and stained with anti-Flag antibody and ALEXA anti-mouse and nuclear DAPI staining, and then examined in a confocal microscope. Flag-proteins are shown in green and nuclei in blue.

### Degradation of xlAZIN2 in HEK293T cells

The half live of xlAZIN2 was calculated by measuring the decay in both enzymatic activity and protein content (estimated by western-blotting), after inhibition of protein synthesis by cycloheximide treatment. Fig 7A shows that xlAZIN2 is a short-lived protein (t_1/2_~ 34 min) with a metabolic turn-over higher than that of xlODC1 (t_1/2_~ 136 min), under the same analytical conditions (Fig 7B). In addition, the great reduction in the degradative rate elicited by treatment with MG132, a potent inhibitor of proteasomal degradation, shown by Fig 7C, suggests that xlAZIN2 can be degraded by the mammalian proteasome in a similar way to that of their mammalian orthologues.

**Fig 7.**
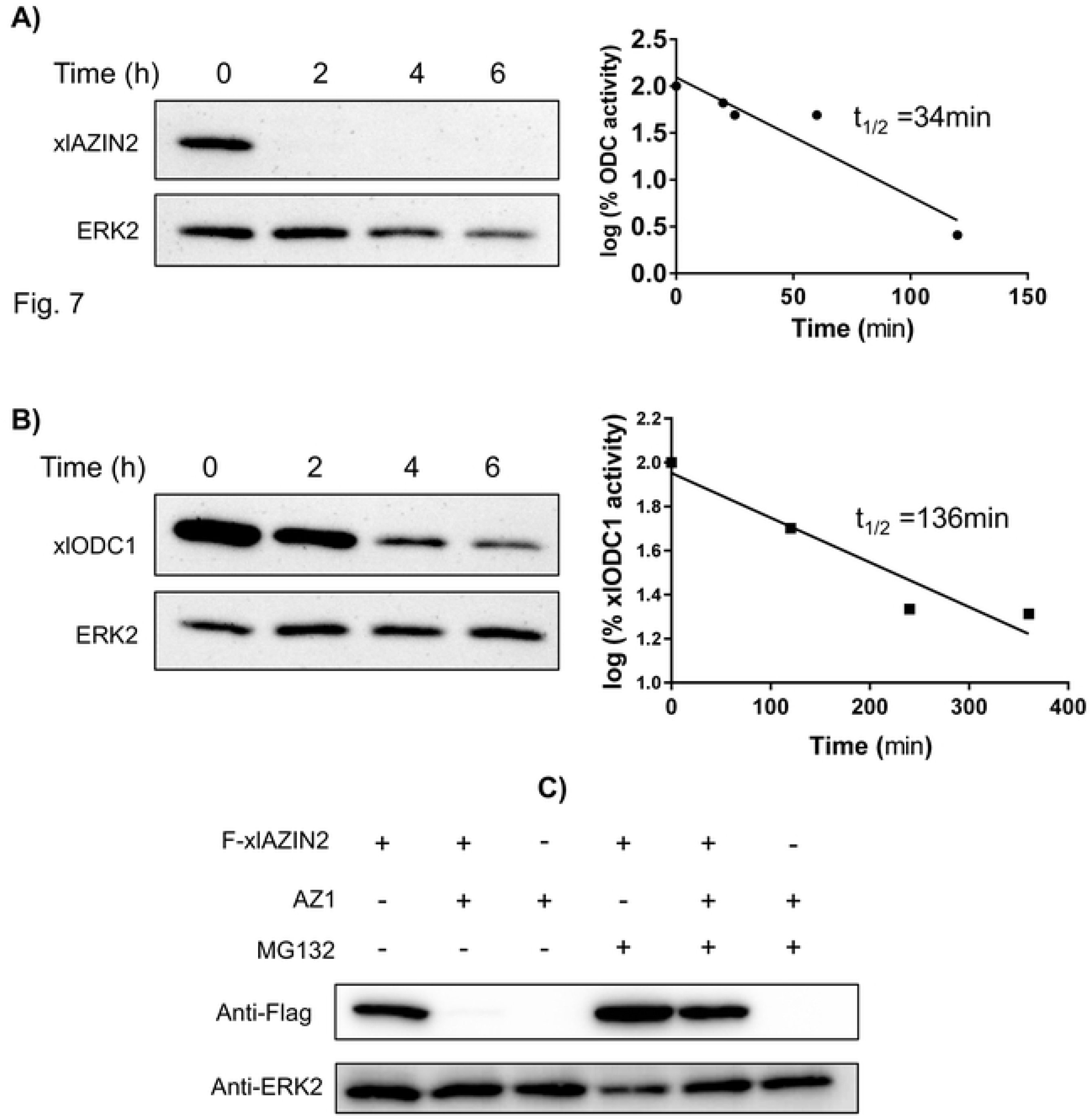
Protein stability of xlAZIN2 and xlODC1 in transfected cells. After 16 h of transfection, either with xlAzin2 or xlOdc1, cells were incubated with 200 μM cycloheximide (CHX), harvested at the indicated times, and lysed in buffer containing a protease inhibitor cocktail. (A) Left: Western blot analysis of xlAZIN2 protein using the anti-Flag antibody; right: decay of ODC activity. (B) Similar experiments with xlODC1. Half-lives of xlAZIN2 and xlODC1 in the transfected cells were calculated by linear regression analysis (GraphPad software). (C) HEK293T cells transfected with xlAZIN2 or xlAZIN2+AZ1were incubated for 5 h with or without the proteasomal inhibitor MG132 (50 μM). xlAZIN2 protein was determined as in (A). ERK2 was used as a loading control.

Since it is very well known that the last 37 amino acid residues of the carboxyl terminus of mammalian ODC play a relevant role in its rapid intracellular degradation [50, 51], we decided to analyze the relevance of this C-terminal region in the amphibian protein on the degradation of xlAZIN2. For that purpose, we generated two mutated versions of xlAZIN2 and studied the influence of the different antizymes (AZ1, AZ2, and AZ3) in the degradation of wild type xlAZIN2, and in that of its C-terminal mutant forms, in the HEK293T-transfected cells. The first xlAZIN2 mutant was a truncated form in which the last 21 amino acid residues of the C-term of xlAZIN2 were deleted (xlAZIN2-ΔC). This deleted sequence (CGWEISDSLCFTRTFAATSII) has a poor homology (14%) with the corresponding one in mODC (CAQESGMDRHPAACASARINV). The second mutant was a quimeric protein (xlAZIN2-mAZIN2) in which the mentioned C-terminal sequence in xlAZIN2 was substituted by the corresponding C-terminal region of mAZIN2 (CGWEITDTLCVGPVFTPASIM). Fig 8A shows that, whereas AZ1, as earlier shown, increased the degradation of xlAZIN2, AZ2 and AZ3 did not stimulate the degradation of this protein. On the opposite, the truncation of the C-terminal region of the protein prevented its AZ1-dependent degradation, indicating that the 21 amino acid residues of the C-terminal region of xlAZIN2, as in the case of mODC, play a relevant role in the degradative process. Again, as in the case of xlAZIN2, the stability of the truncated protein was not significantly affected by any of the other two antizymes (Fig 8B). Moreover, as shown in Fig 8C, the quimeric protein xlAZIN2-mAZIN2 showed against antizymes a behavior similar to that found for xlAZIN2. This indicated that the substitution of the C-terminal of xlAZIN2 by the corresponding region from mAZIN2, did not protect this quimeric protein from the antizyme-induced degradation. Figs 9A and 9B also show that the deletion of the C-terminal region of xlAZIN2 prevented its rapid degradation. The fact that degradation of the quimeric protein xlAZIN2-mAZIN2 was decreased by MG132 (Fig 9C), as already shown by the wild type protein (Fig 7C), suggested that the proteasome participates in the degradation of both proteins.

**Fig 8.**
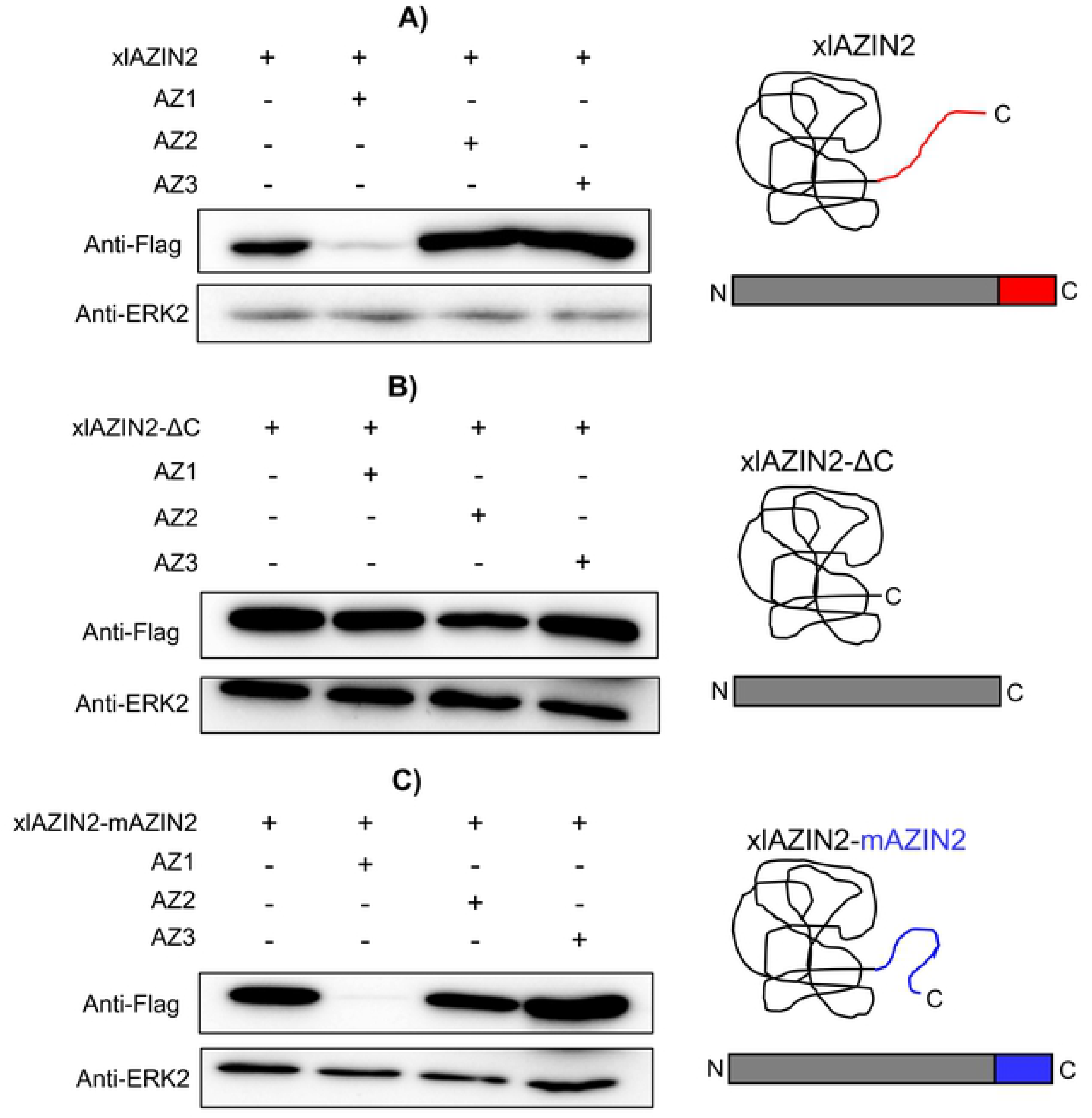
Influence of the C-terminal region of xlAZIN2 in the degradative process induced by AZs. HEK293T cells were transfected with: (A) xlAZIN2, (B) xlAZIN2 lacking the 21 C-terminal residues (xlAZIN2-ΔC) or (C) with a construct coding for a quimeric protein with the substitution of the 21 C-terminal residues of xlAZIN2 by the C-terminal segment of mouse AZIN2 (xlAZIN2-mAZIN2). In parallel, each one of the constructs was co-transfected with members of the AZ family (AZ1, AZ2, and AZ3). Western-blots were probed with anti-Flag antibody. On the right side, schematic representations of xlAZIN2 and the two mutated proteins.

**Fig 9.**
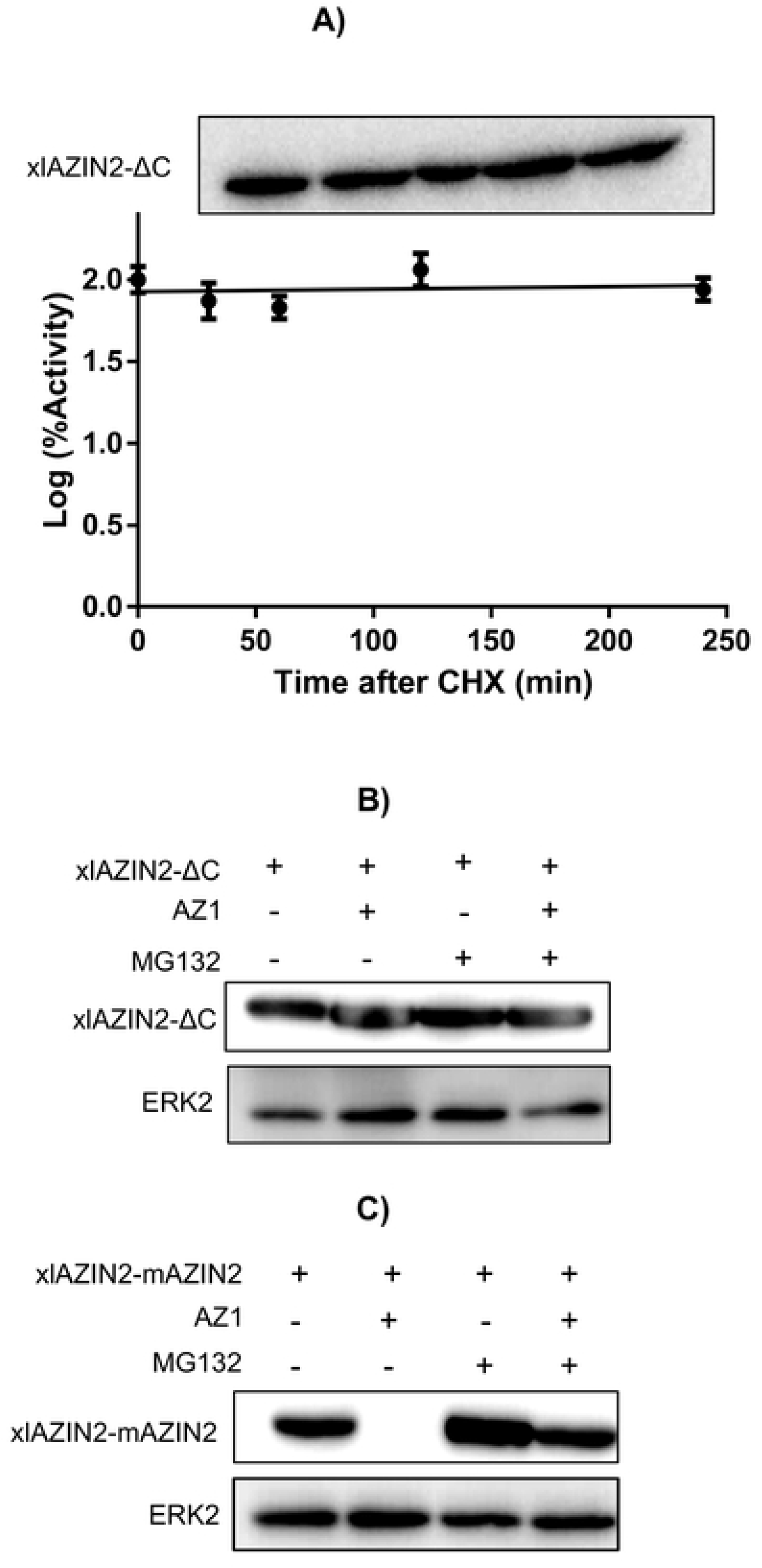
Protein stability of the mutated forms of xlAZIN2. (A) After 16 h of transfection with xlAZIN2-ΔC, cells were incubated with 200 μM cycloheximide (CHX), harvested at the indicated times, and lysed in buffer containing a protease inhibitor cocktail. Top: western blot analysis of xlAZIN2-ΔC at different times after CHX addition; bottom: changes in ODC activity after CHX treatment. (B) Influence of the proteasomal inhibitor MG132 (50 μM) on the effect of AZ1 on xlAZIN2-ΔC protein in HEK293T transfected cells. (C) Influence of the proteasomal inhibitor MG132 (50 μM) on the effect of AZ1 on xlAZIN2-mAZIN2 protein in HEK293T transfected cells.

## Discussion

Our results clearly indicate that xlAZIN2 is devoid of antizyme inhibitory capacity, since it was unable to rescue ODC from the negative effect of AZ1 (Fig 5A). In addition, AZ1 did not protect xlAZIN2 from degradation (Figs 5B and 7C), contrary to what was reported for mAZIN1 and mAZIN2 [26, 37].Unexpectedly, AZ1 accelerated the degradation of xlAZIN2 by the proteasome (Fig 7C), as it was also observed for xlODC1 (Fig 5A), and as early described for mammalian ODCs [14, 52].

On the contrary, our findings unambiguously demonstrated that xlAZIN2 was able to decarboxylate not only L-ornithine but also L-lysine, producing the diamines putrescine and cadaverine, respectively (Figs 4A and 4B). It was also clear that in the cultured cells transfected with either xlODC1 or mODC, cadaverine was also produced but in a lesser amount than putrescine (Figs 4C and 4D). These results are in agreement with early reports that showed that ODC from rodent tissues was able to decarboxylate both amino acids, although L-lysine less efficiently than L-ornithine [49]. The comparison of the kinetic parameters of xlAZIN2 with those of xlODC1 showed that the affinity of xlAZIN2 for lysine is about 30-fold higher than that of xlODC1, whereas the opposite was evident for ornithine. All these results reveal than in *Xenopus laevis* there are two related genes (xlODC1 and xlAZIN2) coding for enzymes able to decarboxylate both amino acids ornithine and lysine. Whereas the function of xlODC1 appears to be related with the formation of putrescine, and therefore in consonance with that of mammalian ODCs, our data suggest that it is very likely that the main role of xlAZIN2could be concerned with the synthesis of cadaverine. Although some studies revealed the presence of cadaverine in several amphibian tissues [53, 54], including those adult *Xenopus laevis* during limb regeneration [55], the physiological function of this diamine is mostly unknown. Taking into consideration that the protein sequence of xlAZIN2 is identical to that reported for xlODC2 [36], it can be assumed that xlODC2 may have lysine decarboxylase activity, although it should be noted that no enzymatic activity for xlODC2 was measured in the mentioned report. Interestingly, it was also reported that xlODC1 and xlODC2 showed different expression patterns during *Xenopus laevis* embryo development [36]. The specific regional and temporal expression of xlODC2 during specific stages of *Xenopus* embryo development [36], associated to the mentioned lysine decarboxylase activity of xlODC2, suggest that cadaverine may have some role during *Xenopus* embryogenesis, different to that of putrescine. This possibility could also explain the reason for the existence of two apparently similar ODC decarboxylases in *Xenopus*.

According to our results, the two *Xenopus* enzymes xlODC1 and xlODC2/xlAZIN2 expressed in mammalian cells share several properties with mouse ODC, such as their cytosolic localization, short half-lives, and AZ1-stimulated degradation by the proteasome. On the other hand, xlODC2/xlAZIN2 differs from mAZIN2 in that the murine protein lacks decarboxylase activity and is located in vesicular-like structures, and that AZ1 protects mAZIN2 from degradation [37, 56, 57]. The mechanisms by which AZs exert opposite effects on the protein stability of ODC and AZINs are not completely understood. Different studies have demonstrated that in ODC there are two regions participating in its rapid turn-over [revised in 58]. The first region encompasses amino acid residues 117-140 needed for AZ binding (AZBE region) [52]. The second is the C-terminal region in mammalian ODC [59–61] or the N-terminal region of yeast ODC [31]. Interestingly, our results showed that the deletion of the 21 amino acid residues of the C-terminal region of xlODC2/xlAZIN2 made the truncated protein more stable and resistant to AZ1-induced degradation by the proteasome. This result is in agreement with early reports showing that the truncation of the carboxyl-terminal segment of mouse ODC prevented its rapid intracellular degradation [50, 59]. However, the substitution of this C-terminal region in xlAZIN2 by the corresponding one of mAZIN2 did not protect it from AZ1-induced degradation, despite being known that the degradation of mAZIN2 is not stimulated by binding to AZ1 [27]. Taking into consideration the low sequence homology between the 21 amino acid C-terminal tail of xlAZIN2 or that of the quimeric protein with that of mODC (14% and 9%, respectively, as shown in S1 Table), our results support the contention that different C-terminal amino acid sequences may lead to the interaction of these ODC homologous proteins with the proteasome. According to current views, an unstructured terminal domain can be absolutely essential as the initiation site for protein degradation [62, 63]. As shown here, in the case of xODC homologues, different C-tail sequences can accomplish this requirement. Apart from the implication of this terminal protein segment (S2) in the initial infiltration of the protein in the proteolytic chamber of the proteasome, recent studies based on structural analyses have proposed that the interaction of ODC-AZ complex with the proteasome requires the exposure of the ODC residues 391-420 [19]. However, the specific role of the different amino acid residues within this pre-terminal sequence (S1) on the interaction with the proteasome is still unknown. As shown in S1 Table, it is clear that the homology of the S1 segment of xlODC1 and xlAZIN2/ODC2, the two amphibian proteins induced to be degraded by AZ1, with that of mODC is higher than those calculated for xlAZIN1 or mouse AZINs, proteins whose degradation is not stimulated by AZ1. This finding supports early conclusions based on structural studies that claimed for the relevance of the 391-420 ODC region for interacting with the proteasome [19]. The existence of the invariable sequence FNGFQ in the S1 segments of mODC, xlODC1 and xlAZIN2/ODC2 (S2 Fig), that according to the above-mentioned structural study forms a short helical turn, suggests that this part of the S1 terminal region may be critical for recognition by the proteasome. If this is the case, the alteration of this sequence in the AZINs could make these proteins resistant to the degradative stimulatory action of AZs.

Collectively, our study demonstrates, firstly, that xlAZIN2, although having a gene structure similar to those of mammalian AZIN2s, is not really an antizyme inhibitor, but an authentic decarboxylase with preference for L-lysine as substrate. According to this, the name of xlLDC (or xlODC2), instead of xlAZIN2, should be used. Secondly, our results also extend the previous knowledge on the influence of AZs on degradative aspects, from mammalian ODCs to non-mammalian ODCs different from yeast or trypanosomal ODCs. Our findings support the hypothesis that in the C-terminal region of *Xenopus* ODCs the last 21 amino acid tail is required for antizyme-stimulated degradation of the enzyme, and suggest that the sequence FNGFQ encompassing residues 396~400 may be relevant for the interaction of mammalian and amphibian ODCs with the proteasome.

## Supporting information

**S1 Fig. Comparison of the amino acid sequences of *Xenopus laevis* AZIN2 (xlAZIN2) and *Xenopus tropicalis* AZIN2 (xtAZIN2) using ClustalW program for multiple sequence alignment**. Asterisks represent amino acid identity; colon and dots represent amino acid similarity between the proteins. Grey background indicates amino acid residues associated with the catalytic activity of mODC that are conserved in the *Xenopus* homologues.

**S2 Fig. Sequences of the C-terminal region of mODC and its paralogues and *Xenopus laevis* orthologue proteins.** (A) Scheme of the C-terminal region of mODC, where C represent the ~70 amino acid residues, and S1 and S2 the two subregions that may be important for proteasomal degradation of ODC induced by AZ1. (B) Detailed sequence of the C-terminal region of mODC and its different paralogues and orthologues. Sequences corresponding to S1 (residues 391-420) and S2 (residues 441-461) are underlined.

**S3 Fig. Influence of AZ1 on protein levels of xlAZIN1**. HEK293T cells were transfected with xlAZIN1-Flag alone or in combination with AZ1. Western-blots were probed with anti-Flag and anti-ERK2 antibodies.

**S1 Table. Sequence identity between the C-terminal region of mODC and those of its *Xenopus laevis* homologues**. C: terminal region from residues 391 to 461 in mODC; S1 and S2 are two subregions that may be important for ODC proteasomal degradation induced by AZ1; S1: residues 391-423; S2: residues 441-461. (See S2 Fig).

